# RNA-binding protein RBM3 negatively regulates innate lymphoid cells (ILCs) and lung inflammation

**DOI:** 10.1101/2020.07.27.223958

**Authors:** Jana H. Badrani, Michael Amadeo, Kellen Cavagnero, Luay H. Naji, Sean J. Lund, Anthea Leng, Lee Lacasa, Allyssa Strohm, Hyojoung Kim, Rachel E. Baum, Naseem Khorram, Monalisa Mondal, Grégory Seumois, Julie Pilotte, Peter W. Vanderklish, Taylor A. Doherty

## Abstract

Innate lymphoid cells (ILCs) promote lung inflammation through cytokine production in diseases such as asthma. RNA-binding proteins (RBPs) are critical post-transcriptional regulators of cellular function, including inflammatory responses, though the role of RBPs in innate lymphoid cells is unknown. Here, we demonstrate that RNA-binding motif 3 (RBM3) is one of the most highly expressed RBPs in Thy1.2+ lung ILCs after fungal allergen challenge and is further induced by epithelial cytokines TSLP and IL-33 in both human and mouse ILCs. Single (*rbm3*−/−) and double (*rbm3*−/−*rag2*−/−) knockout mice exposed via the airway to the asthma-associated fungal allergen *Alternaria alternata* displayed increases in eosinophilic lung inflammation and ILC activation compared to control mice. In addition to increased Th2 cytokine production, *rbm3*−/− ILCs produced elevated IL-17A. The negative regulation by RBM3 in ILC responses was direct as purified *rbm3*−/− ILCs were hyperinflammatory *in vitro* and *in vivo* after stimulation with IL-33. Transcriptomic analysis by RNA-sequencing of *rbm3*−/− lung ILCs showed increased type 2 and 17 cytokines as well as global expression differences in critical cytokines, receptors, transcription factors, and survival transcripts compared with WT ILCs. Intriguingly, these transcript changes did not correlate with the presence of AU-rich elements (AREs), which RBM3 is known to bind. Thus, regulation of ILC responses by RNA-binding proteins offers novel mechanistic insight into lung ILC biology and ILC-driven inflammatory diseases.

## Introduction

Lung innate lymphoid cells (ILCs) are critical players in inflammatory diseases including asthma, viral and helminth infections, and pulmonary fibrosis^1, 2, 3, 4^. Once activated by epithelial cytokines and lipid mediators, group 2 innate lymphoid cells (ILC2s) produce Th2 cytokines IL-4, IL-5, IL-9, and IL-13^5, 6^. These cytokines are major contributors to the characteristics of asthma such as airway inflammation, hyperresponsiveness, and remodeling. IL-13 promotes mucus production and airway hyperresponsiveness, IL-4 regulates IgE synthesis and Th2 cell differentiation, and IL-5 controls the survival and activation of eosinophils^7, 8^. In addition to conventional ILC2 contributions to asthma, there is also evidence that IL-17A production by “inflammatory” ILC2s (iILC2s or ILC2-17s), as well as ILC3s, promotes lung inflammatory responses in asthma models^2, 9, 10^. Since the discovery of ILCs, significant insight regarding mechanisms of activation of ILCs has been gained, while mechanisms of suppression of lung ILC responses are less understood but may occur through individual cytokines^11, 12^, lipid mediators^13, 14^, and post-transcriptional events that regulate the synthesis of these and other factors. Overall, our understanding of novel mechanisms that globally suppress both type 2 and 17 cytokine production in lung ILCs is limited.

Regulation of immune cell responses occurs through gene expression as well as post-transcriptional and post-translational pathways. RNA-binding proteins (RBPs) regulate cellular responses via stabilization or degradation of mRNA transcripts, their spicing and transport, as well as effects on miRNA processing^15^. The RBP human antigen R (HuR), for instance, stabilizes target mRNAs including GATA3 transcripts in CD4+ Th2 cells through binding to AU-rich element (ARE) sequences in the 3’UTR ^16^. In contrast, the RNA binding protein tristetraprolin (TTP/zfp36) negatively regulates inflammation through destabilization of ARE-bearing mRNAs^17^. Importantly, AREs are present in Th2-cytokine transcripts including IL-4, IL-5, IL-13, and are thus potential posttranscriptional regulatory targets for RBPs^18^. Despite known roles for some RBPs in immune cell activation, the role of RBPs in ILC responses has not been reported.

Utilizing RNA sequencing from purified lung ILC subsets based on CD127 (IL-7R) and ST2 (IL-33R) from fungal allergen-challenged mice, we assessed levels of ILC RBP transcripts. RNA-binding motif 3 (*RBM3*) gene was one of the most highly expressed RBPs in ILCs, with a higher expression level than *Elavl1*, which encodes for HuR. RBM3 is a cold shock protein that has been demonstrated to enhance the stability and translation of COX-2, IL-8, and VEGF mRNA^19^. Further, RBM3 promotes translation ^20^ and the biogenesis of a broad set of microRNAs 15, including miR-142–5p and miR-143 which are temperature-sensitive microRNAs implicated in the fever response^21^. RBM3 also interacts directly with the RBP HuR^19^. Here, using fungal allergen- and IL-33-driven lung inflammation models with *rbm3*−/− mice, as well as *in vitro* studies and transcriptomic analysis, we show that RBM3 negatively regulates lung ILC type 2 and 17 cytokine responses which are accompanied by relevant global ILC transcriptomic changes. Contrary to previous reports showing stabilization of cytokine transcripts by RBM3 ^22^, the effects of RBM3 on cytokine responses and transcriptomic changes did not correlate with the presence of ARE’s in affected transcripts. Our data define a novel role for RBM3 in suppression of ILC activation and evidence that RBPs are key regulators of immune responses.

## Results

### RBM3 is highly expressed among RNA-binding protein (RBP) mRNAs in activated ILCs

CD127 (IL-7R) and ST2 (IL-33R) are established ILC2 surface markers, however, we have recently shown that single negative or double negative “unconventional” subpopulations for these markers include heterogenous ILC populations that express GATA-3 and produce type 2 cytokines.^23^ Upon airway challenge with the asthma-associated fungal allergen *Alternaria alternata*, we showed that all four Lin-Thy1.2+ subpopulations (CD127+ST2+, CD127+ST2-, CD127-ST2+, and CD127-ST2-) are activated and therefore we did not limit our current analysis to double positive CD127+ST2+ ILCs. Lin-Thy1.2+ lymphocyte subsets from WT mice challenged over 10 days with 50μg *Alternaria* extract were FACS purified based on expression of CD127 and ST2 surface markers (Supplementary Fig. 1a) and subsequently processed for RNA-seq library preparation and sequencing (see methods). Gene expression analysis of the 4 populations revealed that 207 RNA-binding proteins (RBPs) were expressed in the ILC subpopulations (Supplementary Fig. 1b). Of the top 25 differentially expressed RBPs (Fig. 1a), *zfp36*, which encodes for tristetraprolin (TTP), was the most highly expressed RBP. Previous studies have demonstrated that *Zfp36*−/− mice develop spontaneous prominent inflammation and severe autoimmune disease^24^. As we were interested in studying novel RBP influences on ILC biology using *in vivo* asthma models, we focused on a potential role for RBM3 given that these mice are viable and not known to spontaneously develop immune disease^25, 26^. After *zfp36, Rbm3* was the next highest RBP with transcript levels significantly upregulated (400 transcripts per million) by allergen exposure, four-fold higher than *elavl1* that encodes for HuR (Fig. 1b), which has been shown to promote GATA-3 expression and type 2 cytokines in T cells^16, 27^. In qPCR studies, we found that sorted Thy1.2+ lung ILCs from *Alternaria*-challenged mice expressed *rbm3* mRNA at levels on par with *il5* mRNA though less than *gata3*, *rora*, and *il13* (Fig. 1c). Thus, of 207 RBP transcripts from *in-vivo* activated lung ILCs, RBM3 was highly expressed and thus might regulate ILC function and lung inflammation.

**Fig. 1:**
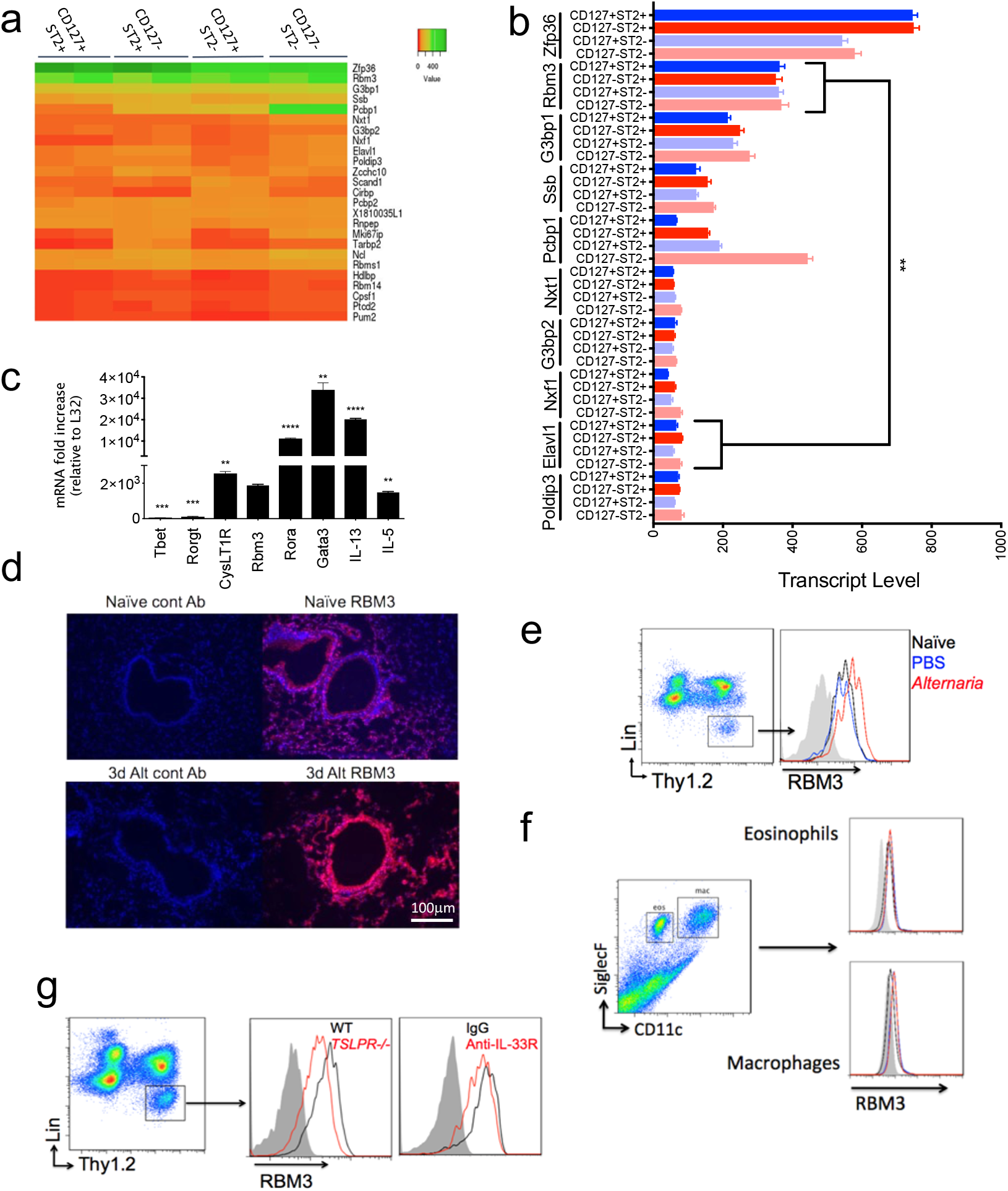
RBM3 is highly expressed in mouse ILCs and is induced by *Alternaria*, IL-33 and TSLP. (a) Wild-type mice were challenged four times with 50μg *Alternaria* over 10 days and four lung ILC subsets (CD127+ST2, CD127-ST2+, CD127+ST2-, CD127-ST2-) were sorted and collected for RNA-seq. (a) Heatmap showing the relative expression of the top 25 highly expressed RBPs in the four ILC subtypes. RBPs involved in homeostatic cell cycle or splicing were excluded. Centroid linkage and Manhattan clustering used. (b) Absolute transcript levels of the top 10 RBPs in the four ILC subtypes. (c) RBM3 mRNA levels of ILCs in comparison to several traditional ILC2 mRNA (*IL-5, IL-13, Gata-3, Rora,* and *CysLT1R*) and non-ILC2 mRNAs (*Rorgt* and *Tbet*). Data are from triplicate samples. Values compared with Rbm3. (d) Immunofluorescence of RBM3 expression in airways of naïve and *Alternaria* challenged mice. Cell nucleus was stained with DAPI. Images taken from at least 5 airways of at least 3 mice. (e) RBM3 expression in ILCs from naive, PBS challenged, or *Alternaria* challenged lung. Black = Naive, Blue = PBS challenged, Red = *Alternaria* challenged (24 hour). (f) RBM3 expression in macrophages and eosinophils from naive, PBS challenged, or *Alternaria* challenged lung. Black = naive, blue = PBS challenged, red = *Alternaria* challenged (24 hour). (g) RBM3 expression on lung ILC2s of *tslpr*−/− mice and mice treated with anti-IL-33R. Data representative of at least 4 mice (paired) per group. Mice were challenged with *Alternaria* four times. Grey = isotype control, black = wild-type or IgG treated, red = *tslpr*−/− or anti-IL-33R treated. * p < 0.05, ** p < 0.01, *** p < 0.001, Unpaired t-test.

### Alternaria, IL-33, and TSLP increase lung and ILC RBM3 expression

In order to determine whether RBM3 is upregulated at the protein level during type 2 inflammation, *Alternaria*-challenged mice were assessed for RBM3 levels by immunostaining. Increased expression of RBM3 was detected by immunofluorescence in *Alternaria*-challenged lungs compared to naïve controls (Fig. 1d). RBM3 expression was visualized in epithelial as well as subepithelial cells which was further induced by challenge with *Alternaria*. As ILCs regulate innate lung responses *to Alternaria*^*28*^, we investigated changes in RBM3 levels in lung ILCs. Lin-Thy1.2+ ILCs from the lungs of challenged mice demonstrated increased RBM3 expression by intracellular flow cytometry when compared to naïve and PBS-challenged controls (Fig. 1e). Interestingly, eosinophils and macrophages analyzed from challenged mice did not show increases in RBM3 expression after *Alternaria* challenge (Fig. 1f). As IL-33 and TSLP are critical epithelial cytokines in type 2 inflammation that regulate ILC2 responses, we assessed levels of RBM3 in *tslpr*−/− mice and WT mice treated with IL-33 blocking antibody. RBM3 expression in *tslpr*−/− mice and WT mice receiving anti-IL-33R was reduced compared with controls (Fig. 1g). Overall, these data suggest that RBM3 is induced in mouse ILCs during type 2 lung inflammation and regulated in part by TSLP and IL-33.

We next assessed whether human ILC2s expressed RBM3 and if TSLP and IL-33 regulated RBM3 expression similar to mouse ILCs. Lin-CRTH2+ ILC2s from human nasal polyp expressed RBM3 suggesting pathologic human airway tissue ILC2s express RBM3 (Fig. 2a). Further, FACS purified human ILC2s stimulated with a combination of TSLP and IL-33 demonstrated significantly increased RBM3 immunostaining after 24 and 72 hours compared with control cells (Fig. 2b). These findings were consistent with increased intracellular RBM3 staining in ILC2s from human PBMCs stimulated with IL-33 or TSLP alone or in combination (Fig. 2c). Thus, activation of ILC2s induced increased RBM3 expression in mouse and humans, suggesting a potential regulatory role of RBM3 in ILC function.

**Fig. 2:**
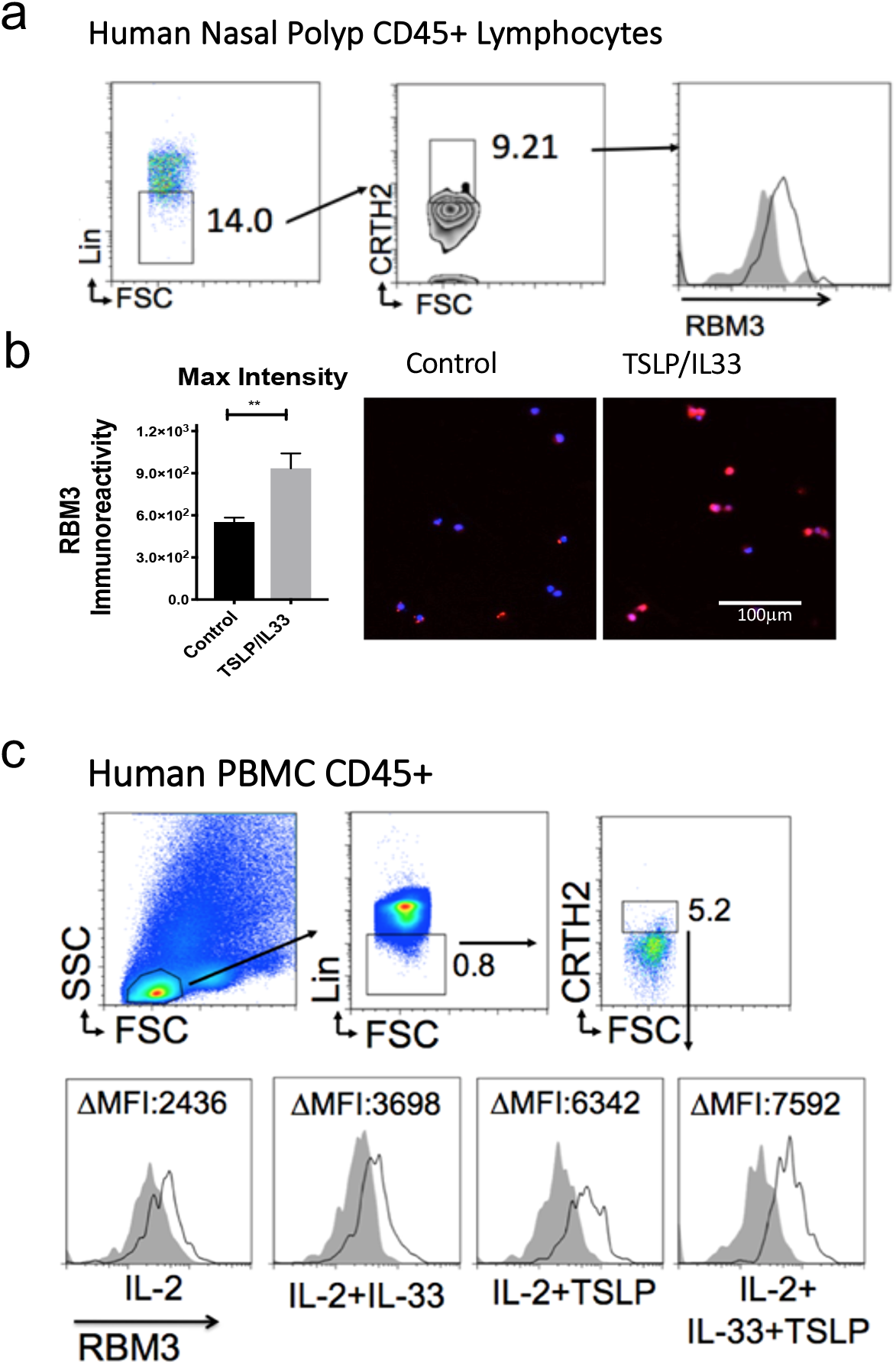
Human ILC2s express RBM3 which is induced by TSLP and IL-33. (a) RBM3 expression in Lineage-CRTH2+ ILC2s from human nasal polyp. (b) Human ILC2s were purified by FACS and cultured and treated with TSLP and IL-33 before being fixed with 4% PFA and processed for immunocytochemistry using a RBM3 antibody (at 1:2000) and a Cy3 secondary. Cells were also stained for DAPI. Images were taken at 20X. Immuno-positive elements were captured and analyzed by thresholding the intensity histogram in the Cy3 channel at 100. Max intensity after 24 hours of stimulation were quantified. Data for objects of 50-175 pixels were included. Immunostaining shown after 72 hours of stimulation. (c) RBM3 expression on ILC2s from isolated human PBMCs. ILC2s were cultured with IL-2, IL-33, TSLP, or a combination. ** p < 0.01, Mann-whitney.

### RBM3 suppresses Alternaria-induced type 2 lung inflammation

To assess the potential role of RBM3 in ILC function and lung inflammation, we performed *in-vivo* asthma model experiments with *rbm3*−/− mice. We analyzed lineage-negative Thy1.2+ lung ILCs from naive *rbm3*−/− mice and found no significant changes in the absolute or relative number of ILCs between naïve WT and *rbm3*−/− mice (Supplementary Fig. 2a, b). Naive *rbm3*−/− ILCs had similar surface marker expression of common ILC2 surface markers when compared to WT ILC2s (Supplementary Fig. 2c) and naïve *rbm3*−/− mice also showed similar levels of BAL and lung eosinophils as WT controls (Supplementary Fig. 2d, 2e). Thus, *rbm3*−/− mice are phenotypically similar to wild-type mice in terms of baseline lung eosinophil and ILCs levels.

In order to assess the role of RBM3 *in vivo*, we repetitively challenged WT and *rbm3*−/− mice with *Alternaria*, a fungal allergen that induces potent type 2 lung inflammation and ILC activation^23, 28, 29^. In multiple challenge protocols, we observed increased eosinophilia in *rbm3*−/− mice, as early as 24 hours in both BAL and lung (data not shown). *rbm3*−/− mice challenged with *Alternaria* three times over 7 days showed significant increases in BAL and lung eosinophils (Siglec-F+ CD11c-cells) and neutrophils (Siglec-F-GR-1+ cells) compared to WT mice (Fig. 3a, b). Similarly, *rbm3*−/− mice receiving 4 challenges over 10 days also showed increased eosinophils and neutrophils in BAL and lung (Fig. 3c, d), as well as increased BAL IL-5 and IL-13 (Fig. 3e). Further, lung sections stained for H&E and PAS showed significantly increased peribronchial inflammation and mucus production in *rbm3*−/− mice (Fig. 3f). Thus, in multiple *Alternaria* challenge models, *rbm3*−/− mice exhibited increased lung inflammation when compared to WT mice.

**Fig. 3:**
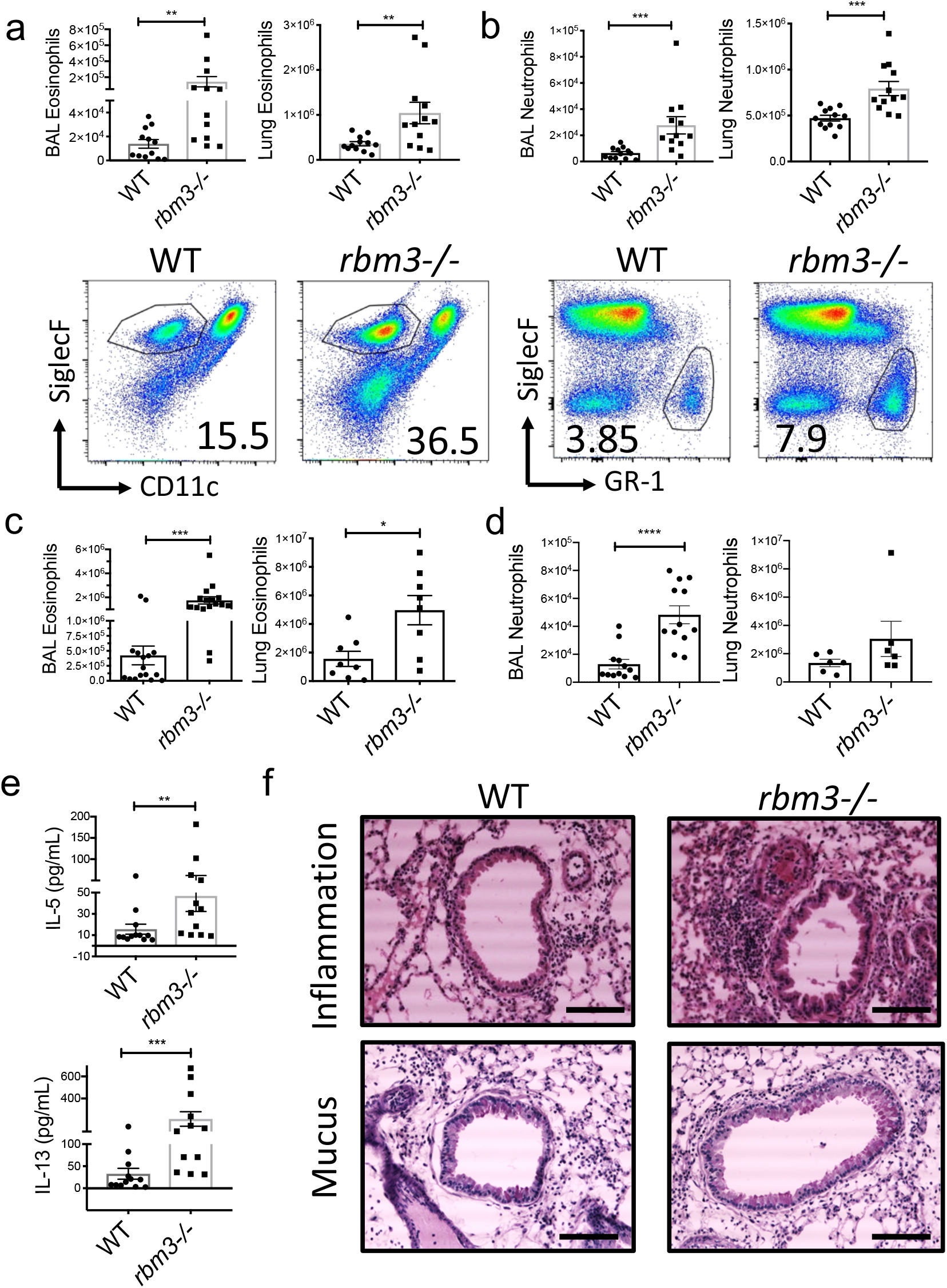
RBM3 suppresses lung inflammation in *Alternaria-* challenged mice. Wild-type and *rbm3*−/− mice were challenged with 10 μg *Alternaria* three times over 7 days. Data shown is from 3 experiments (4 mice per group). (a) Total BAL and lung eosinophils with representative FACS plots of WT and *rbm3*−/− mice. (b) Total BAL and lung neutrophils with representative FACS plots. Mann-Whitney Test. Wild-type and *rbm3*−/− mice were challenged with 20μg and 10μg *Alternaria* over the span of 10 days. Data shown from 4 experiments (4 mice per group). (c) Total BAL and lung eosinophils. Mann-Whitney Test. (d) Total BAL and lung neutrophils. (e) BAL levels of Type 2 cytokines. (f) H&E and PAS lung sections at 20X; scale bar is 100 μm. Airway images representative of 4 mice per group. * p < 0.05, ** p < 0.01, *** p < 0.001, Mann-Whitney Test.

### RBM3 suppresses Alternaria-induced lung ILC responses

We next determined whether ILC activation and proliferation were present in the lungs of *rbm3*−/− mice challenged with *Alternaria*. Lung ILCs identified as CD45+ lineage-negative Thy1.2+ lymphocytes were increased in both number and percent in challenged *rbm3*−/− mice compared with WT mice (Fig. 4a). Further, Ki67+ proliferating ILCs were also increased significantly in *rbm3*−/− mice (Fig. 4b). ILC2s identified by their production of IL-5 and IL-13 were higher in the *rbm3*−/− mice compared with WT mice (Fig. 4c). Interestingly, ILCs from *rbm3*−/− mice demonstrated significantly higher production of IL-17A which may be produced by iILC2s (or ILC2-17s) and/or ILC3s (Fig. 4d)^2, 9, 10^.

**Fig. 4:**
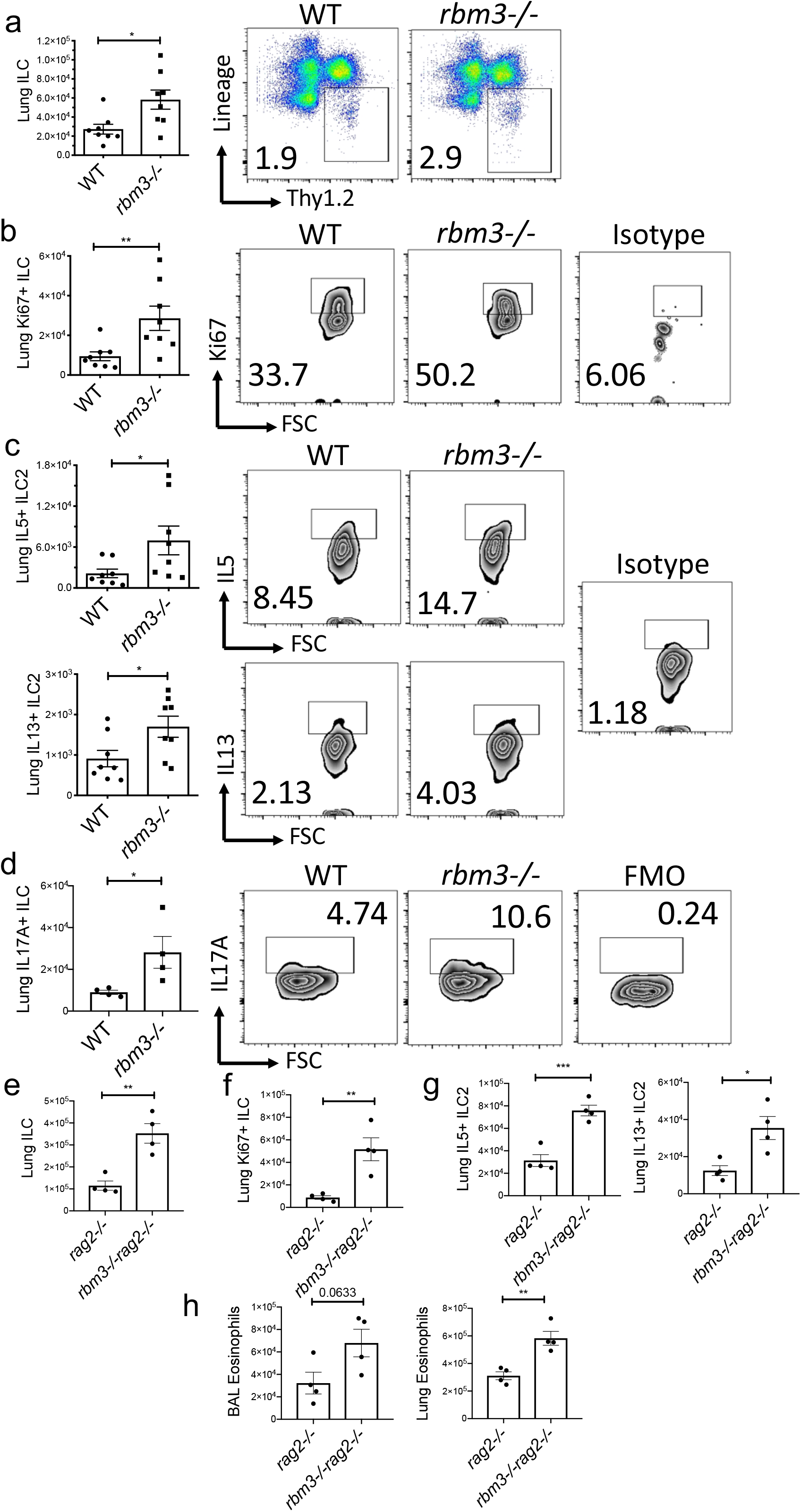
Exaggerated lung ILC type 2 and 17 cytokine production in RBM3−/− mice independent of adaptive immunity. Wild-type and *rbm3*−/− mice were challenged with 20μg and 10μg *Alternaria* over the span of 10 days. Data shown are representative of 4 experiments (4 mice per group). (a) Lung ILC totals and FACS plots of Lin-Thy1.2+ lymphocytes, *p<0.05, Mann-Whitney Test. (b) Totals of Ki-67 expressing lung ILCs. FACS plots of Ki-67 percentages and isotype control, *p<0.05, Mann-Whitney Test. (c) Total IL5 and IL13 expressing lung ILCs. FACS plots of type 2 cytokines percentages and isotype control, *p<0.05, Mann-Whitney Test. Mice were challenged with 25μg *Alternaria* three times over 7 days. Data show from 4 mice per group. (d) Total IL17A expressing ILCs and representative FACS plots of IL17A percentages and FMO control. Unpaired t Test. *rag2*−/− and *rbm3*−/−*rag2*−/− mice were challenged with 20 μg *Alternaria* four times over 10 days. Data representative of 4 mice per group. (e-g) Total number of ILCs, Ki67 expressing ILCs, and type 2 cytokine producing ILC2s. Unpaired t Test. (h) Total BAL and lung eosinophils. Unpaired t Test. * p < 0.05, ** p < 0.01, *** p < 0.001

We also created double deficient *rbm3*−/−*rag2*−/− mice to better assess the role of RBM3 in ILC activation and lung inflammation in the absence of T and B cells. Consistent with the findings in single knockout *rbm3*−/− mice, the *rbm3*−/−*rag2*−/− mice demonstrated significantly higher lung ILCs, Ki-67-expressing ILCs, and type 2 cytokine-producing ILC2s compared to *rag2*−/− mice (Fig. 4e, f, g). Similarly, BAL and lung eosinophils were also significantly increased in the double knockout mice compared to controls (Fig. 4h). Overall, these results show a suppressive role for RBM3 in allergen-induced lung inflammation and ILC activation, even in the absence of adaptive immunity.

### RBM3 directly suppresses IL-33-induced ILC responses in vitro and in vivo

To determine whether RBM3 directly regulated ILC responses, we performed studies with IL-33 which directly activates ILCs *in vitro* and *in vivo*. FACS sorted WT and *rbm3*−/− lung Lin-Thy1.2+ ILCs from *Alternaria-*challenged mice were rested for 48 hours in media alone prior to stimulation. Immediately prior to stimulation, *rbm3*−/− ILC2s produced significantly more IL-5 by ELISA than wild-type ILC2s (Fig. 5a). In contrast, there were no differences in the production of IL-13 prior to stimulation. IL-33 stimulation (15 ng/ml) of *rbm3*−/− ILCs resulted in increased IL-5 and IL-13 compared with wild-type ILCs (Fig. 5b). ILCs lacking RBM3 produced significantly more IL-5 at both doses (15 and 30 ng/ml) of IL-33. Consistent with data in Fig. 4d, ILC IL-17A production was increased in *rbm3*−/− ILCs after 48 hours of rest (Fig. 5c). Unlike the Th2 cytokines, IL-17A was not increased above pre-stimulation conditions with IL-33 stimulation (Fig. 5c). These results demonstrate a direct inhibitory regulatory role of RBM3 in ILC type 2 cytokine production in response to IL-33, as well as RBM3 negative regulation of IL-17A detected *ex vivo*.

**Fig. 5:**
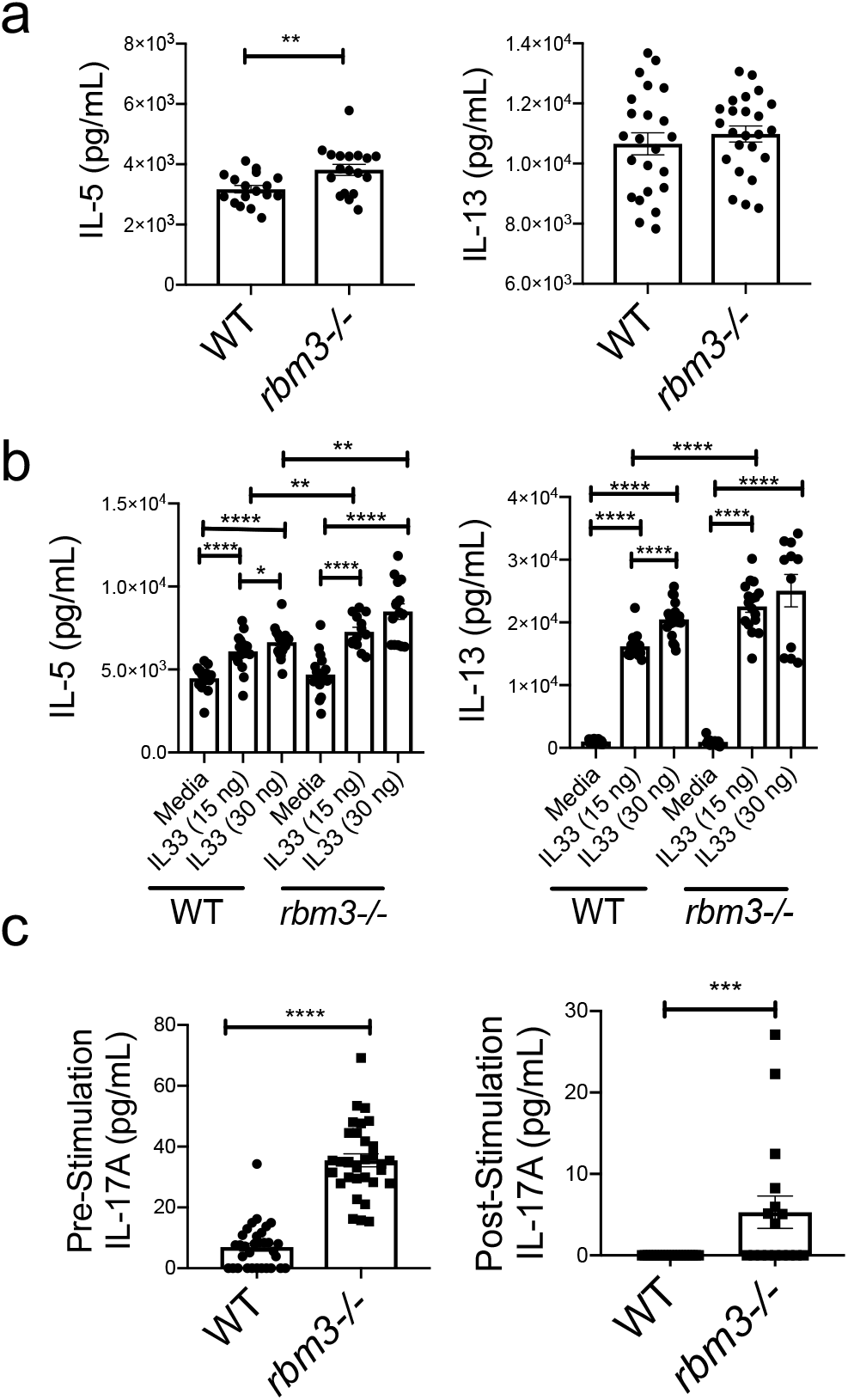
RBM3 directly suppresses lung IL-33-induced ILC activation *in vitro*. WT and *rbm3*−/− mice were intranasally challenged four times with 50 μg *Alternaria* over 10 days. Lin-Thy1.2+ ILCs were FACS sorted from mouse lung and were rested in TCM with 10ng/mL IL-2 and IL-7 for 48 hours prior to stimulation with IL-33. (a) IL-5 and IL-13 concentration by ELISA pre-stimulation. (b) IL-5 and IL-13 concentration post-stimulation with 15 ng and 30 ng IL-33 for 24 hours. (c) IL-17A concentration levels pre-stimulation and post-stimulation with 30n ng IL-33. * p < 0.05, ** p < 0.01, *** p < 0.001, Mann-Whitney Test.

To assess a direct effect of RBM3 function in ILCs *in vivo*, *rbm3*−/− and WT mice were administered IL-33 intranasally which directly activates lung ILC2s^29, 30^. Similar to *Alternaria*-challenged mice, direct challenge with IL-33 led to significantly greater lung Lin-Thy1.2+ ILCs as well as Ki-67+ proliferating ILCs in *rbm3*−/− mice compared to WT controls (Fig. 6a, b). Further, *rbm3*−/− lung cells had significantly higher IL-5 and IL-13 producing ILC2s (Fig. 6c). BAL and lung eosinophilia were also significantly increased in *rbm3*−/− mice (Fig. 6d). A much smaller effect was detected in BAL and lung neutrophil differences after IL-33 administration compared with *Alternaria* models suggesting that this complex allergen has a multitude of inflammatory effects apart from IL-33^31^ (Fig. 6e, Fig. 3). Overall, these findings support that RBM3 suppresses lung ILC activation by IL-33 *in vitro* and *in vivo* in a cell autonomous manner.

**Fig. 6:**
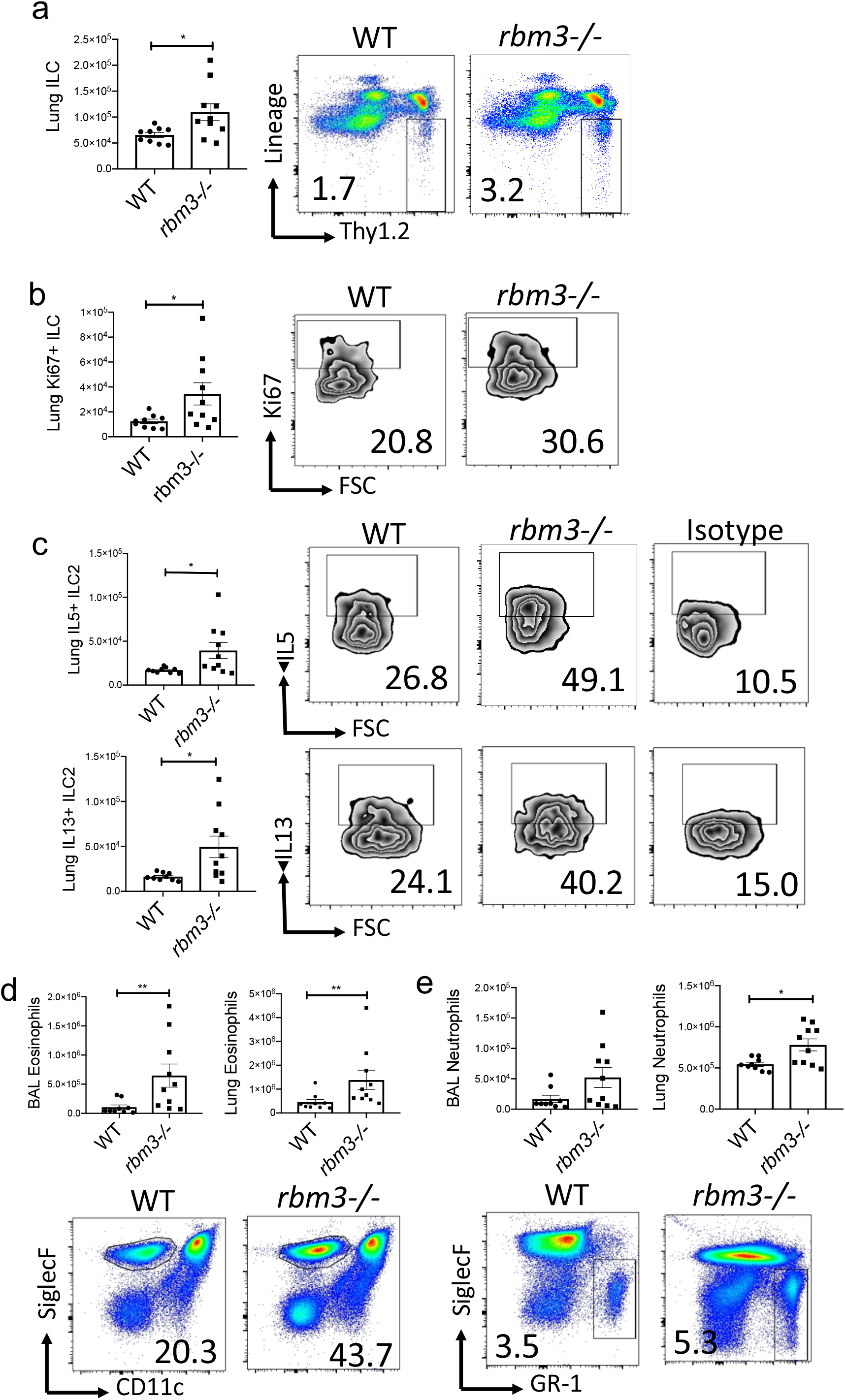
RBM3 directly suppresses lung IL-33-induced ILC activation *in vivo.* WT and *rbm3*−/− mice were intranasally challenged with 10 ng of IL-33 three times over 7 days. (a) Total Lin-Thy1.2+ ILCs and representative FACS plots. (b) Total Ki-67-expressing ILCs and representative FACS plots. (c) Total number of IL5-expressing ILC2s and IL13-expressing ILC2s and representative FACS plots of cytokine levels. Cells were cultured for 3 hours with cell stimulation cocktail prior to staining. (d) Total BAL and lung eosinophils and representative FACS plots of BAL eosinophil levels. (e) Total BAL and lung neutrophils and representative FACS plots of BAL neutrophil levels. Data is from 9-10 mice per group. * p < 0.05, ** p < 0.01, *** p < 0.001, Mann-Whitney Test.

### Activated ILC programs are present in the rbm3−/− ILC transcriptome

We next performed RNA-seq analysis of FACS-sorted Lin-Thy1.2+ ILCs from WT and *rbm3*−/− mice to more broadly understand RBM3 inhibition of lung ILC responses. Gene expression analysis of common ILC transcripts, particularly those related to ILC2s and ILC3s^32, 33^, were more expressed in *rbm3*−/− ILCs than WT controls (Fig. 7a). Principle component analysis of the most variable genes revealed clear transcriptional differences between WT and *rbm3*−/− ILC transcriptomes (Fig. 7b). Level of expression for several cytokine transcripts (*il4*, *il5*, *il6*, *il13*, *areg*, *il17a*, *il17f*, and *il22*) were statistically increased in *rbm3*−/− compared with WT ILCs (Fig. 7c). The ILC1 cytokine *ifng* was not significantly different in *rbm3*−/− ILCs (Fig. 7c) and may reflect specificity to type 2 and 17 ILC responses or overall lack of ILC1 responses in the *Alternaria* model. Further, *rbm3*−/− ILCs had moderate and significant increases in relevant ILC surface markers including *il1rl1, il2rg*, *cysltr1*, *il1rap*, *cd44, klrg1*, and *icos* (Fig. 7d). Interestingly, while type 2 cytokine transcripts were upregulated in *rbm3*−/− ILCs, GATA3 levels were not increased at a transcript or protein level (Fig. 7e, Supplementary Fig. 3c) suggesting other signaling factors are responsible for the increased ILC type 2 cytokine production in *rbm3*−/− ILCs. One such mechanism in *rbm3*−/− ILCs may be through increased *nfactc2* expression that encodes NFAT1 and activates ILC2s via the CysLTR1^34^ to produce Th2 and Th17 cytokines (Fig. 7d, e). Transcription factors involved in the development, differentiation, or activation of ILCs were also upregulated in *rbm3*−/− ILCs, including *tox*^35^, *ets1*^36^, *rora*^37, 38^, *irf4*^39^, and *id2*^40^ (Fig. 7e). ILC ID2 protein expression was also higher in *Alternaria*-challenged *rbm3*−/− mice (Supplementary Fig. 3b). Anti-apoptotic and survival transcripts, including *bcl2*^41^, *cflar*^42^, *bcl2a1d*^43^, and *hbixip*^44, 45^, were moderately or significantly upregulated in *rbm3*−/− ILCs (Fig. 7f), and Bcl2 protein expression was also higher in ILCs from *Alternaria*-challenged *rbm3*−/− mice than WT controls (Supplementary Fig. 3a), suggesting a potential survival benefit to the *rbm3*−/− ILCs. Though RBM3 has been shown to bind to the AU-rich regions of some mRNAs, there were surprisingly no correlations between the number of canonical AUUUA (preferentially bound by RBM3) motifs or total ARE regions and the log fold change of differentially expressed transcripts (Supplementary Fig. 4). Therefore, the AU-rich density of transcripts was not predictive of RBM3 regulation of ILC transcriptome differences.

**Fig. 7:**
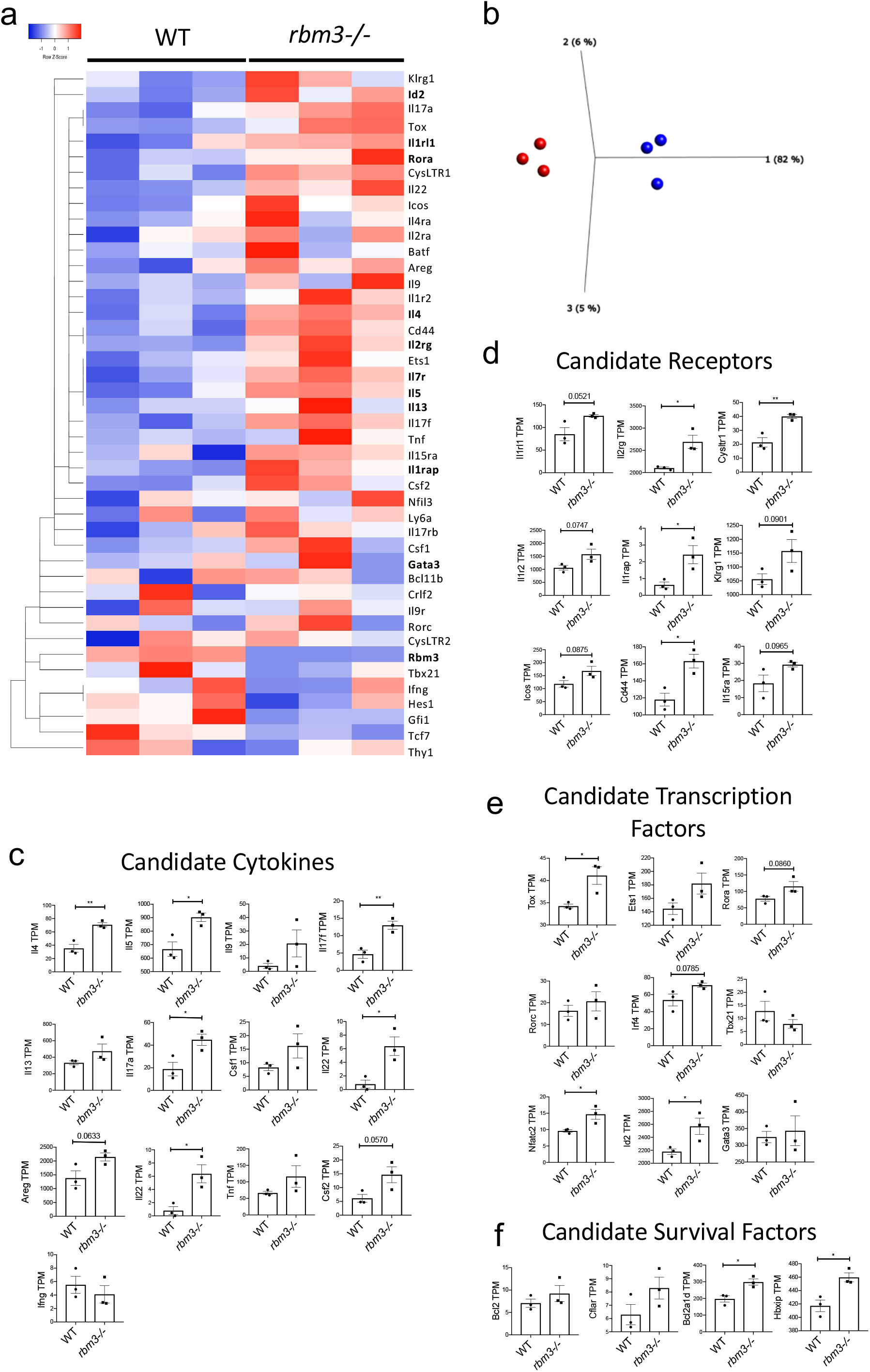
Transcriptome analysis reveals RBM3 control over global ILC activation changes. WT and *rbm3*−/− mice were challenged with 25 μg *Alternaria* three times over 7 days. Lin-Thy1.2+ ILCs were FACS-sorted and bulk RNA-sequenced as per methods. (a) Relative transcript levels of select ILC transcripts. One minus Kendall’s correlation and single linkage used. (b) PCA plot of WT (blue) and *rbm3*−/− (red) samples. (c) TPMs of select cytokine transcripts. (d) TPMs of select surface marker transcripts. (e) TPMs of select transcription factor transcripts. Unpaired t Test. (f) TPMs of select transcripts involved in survival and apoptosis. * p < 0.05, ** p < 0.01, *** p < 0.001, Unpaired t Test.

## Discussion

Innate lymphoid cells including ILC2s have recently emerged as critical contributors to immune diseases including asthma^46, 47^. The majority of the literature thus far has identified soluble factors that activate or inhibit ILC2 function including cytokines and lipid mediators^48^. However, an understanding of novel intracellular mechanisms that more broadly control ILC function might provide important insights into ILC-driven immune diseases. Studies thus far have demonstrated that specific miRNAs activate ILC2s, likely through multiple mechanisms^49, 50^. In this work, we have identified that RNA binding proteins, specifically RBM3, represent a novel aspect of ILC regulation. We demonstrated that RBM3 is a highly expressed RBP in ILC subsets, is induced by epithelial cytokines IL-33 and TSLP, and negatively regulates type 2 and 17 cytokine production by lung ILCs. Further, transcriptomic studies revealed global effects of RBM3 that regulate ILC-relevant cytokines and receptors, transcription factors and survival transcripts.

RBM3 is a 17KD RNA-binding protein with known roles in cellular protection, neural plasticity and oncology^25, 51, 52^. However, very little is understood about the role of RBM3 in inflammation and immunity. One report showed that RBM3 is downregulated in febrile illness and knockdown of RBM3 led to increases in miRNAs that suppress *PGE2*, *IL6*, and *IFNA1*^21^. However, earlier studies showed that RBM3 deficient mice had normal numbers of NK, T, and B cells and had no differences in innate cytokine responses to the TLR9 ligand CpG^26^. We observed that IL-33 and TSLP treatments induced RBM3 expression in vitro, supporting a direct role of signaling events induced by these cytokines in RBM3 induction. While these pathways remain to be elucidated, it is notable that activation of NF-κB pathways promotes RBM3 expression and extracellular IL-33 activates NF-κB pathways^51, 53^. In addition, RBM3 is induced by physiological stresses, including tissue hypoxia and lowered temperatures^52^. Of note, IL-33 has been reported to promote a tumor hypoxic microenvironment, with generation of reactive oxygen species, that could lend toward an indirect induction of RBM3 through local hypoxia^54, 55^. Thus, in vivo there may be both direct and indirect routes to RBM3 induction during inflammatory stress conditions.

Surprisingly, we found that RBM3 suppresses ILC Th2 and Th17 cytokine production as well as ILC proliferation. Furthermore, exacerbation of type 2 responses in *rbm3*−/− mice was independent of changes in total ILC GATA3 levels and lacked correlation with number of ILC AU-rich element (ARE) transcripts. RBM3 has a multitude of complex potential mechanisms that regulate cellular changes during stress including promoting translation rates, the stability of some ARE bearing mRNAs, effects on miRNAs directly or through dicer processing and protein-protein interactions^15, 20, 21, 52^. Interestingly, RBM3 has also been reported to inhibit the p38 MAP kinase pathway which promotes cytokine production by ILC2s in response to IL-33^56, 57, 58^. Thus, removal of RBM3’s inhibition of the p38 pathway in ILCs may result in increased cytokine production in *rbm3*−/− ILCs. It is also possible that RBM3’s effects on ILC cytokine expression involves the microRNA pathway. IL-13 expression and allergic airway inflammation is inhibited by let-7 microRNAs^59^, the biogenesis of which is strongly promoted by RBM3 at the Dicer step^15^. Our studies also demonstrate that RBM3 has no effect on ILC2 numbers and lung eosinophilia under homeostatic naïve conditions but has a clear suppressive role during type 2 inflammatory insult. This may be consistent with RBM3’s role as a “stress-response” protein that strongly exhibits cell protective influences in multiple contexts, including brain and skeletal muscle^52, 60^. Perhaps, limiting of hyperactive ILC responses by RBM3 is an important protective mechanism during lung inflammation.

In this study, we took a broad approach to ILC identification and included lineage-negative Thy1.2+ lymphocytes as ILCs versus specific subsets using conventional surface markers. This is based on our recent work showing that CD127 and ST2 exclude approximately 40% of Th2 cytokine producing ILC2s^23^. Further, several studies have demonstrated significant heterogeneity and plasticity of ILCs (reviewed in^61^). For example, in addition to conventional ILC3s, IL-17 production occurs from other ILC sources including “inflammatory” iILC2s (or ILC2-17s) induced by IL-25, cysteinyl leukotrienes, and notch signaling^9, 10, 62^. In our studies, we did not determine whether the IL-17 production that is enhanced in *rbm3*−/− ILCs is from ILC2-17s or ILC3s. However, our *in vitro* data showed that purified lung Thy1.2+ ILCs produced IL-17 from *Alternaria*-challenged mice and supports that ILC2-17s or iILC2s are the likely source (Fig. 5c). This is based on previous work showing that the vast majority of mouse lung Thy1.2+ ILCs expanded by *Alternaria* airway challenges are activated ILC2s^23, 63, 64^. Therefore, we would expect that ILC2s are largely responsible for the IL-17 production, but we cannot exclude independent populations of ILC3s having some contribution that may be increased in *rbm3*−/− mice.

Our transcriptomic studies suggest global changes in activation of ILCs by RBM3 including differential expression of known ILC cytokines and receptors as well as transcription factor and survival transcripts. Importantly, cytokine transcript data from *rbm3*−/− ILCs supported our *in vivo* findings that RBM3 suppressed ILC Th2 and Th17 cytokine production at a protein level. Multiple receptor transcripts were increased and include *il1rl1* (encoding ST2), *il7r*, *il2rg*, *cysltr1*, and *cd44*. Though *il1rl1* was increased at a transcript level, we did not detect increased ST2 at a protein level by flow cytometry (not shown). Increased expression of the *cysltr1* transcript in *rbm3*−/− ILCs is particularly interesting in combination with increased *nfat2c* (encodes NFAT1) as CysLT1R signaling in ILCs is regulated by NFAT1 to promote increased Th2 and Th17 cytokine production^10, 63, 64, 65^. Thus, RBM3 suppression of the CysLTR1/NFAT1 axis could potentially limit ILC-driven lung inflammation, though it is likely that other RBM3-dependent mechanisms including effects on other transcription and survival factors contribute. Notably, several ILC developmental transcripts were increased in *rbm3*−/− ILCs including *tox*, *ets1*, *rora*, and *id2*. In addition to being ILC developmental factors, Rorα and ETS1 also promote ILC2 cytokine production suggesting that the mature ILC2 cytokine production could be regulated by RBM3 through control of these transcription factors^36, 66^. Despite differences in developmental gene levels in *rbm3*−/− ILCs, we did not detect differences in lung ILCs in naive *rbm3*−/− mice. This may be explained by RBM3 induction in inflammatory settings that then exerts effects on mature ILCs and is dispensable for ILC development. Overall, given the global changes in lung ILC transcriptome regulated by RBM3, it is likely that no one single mechanism is responsible for the *rbm3*−/− ILC phenotype found *in vivo* during lung inflammation. Future avenues of study including miRNA regulation by RBM3 may add to the current studies demonstrating suppression of ILC responses during lung inflammation.

In summary, this work is the first to identify a role for RBM3 – an RNA-binding protein that is highly expressed in activated lung ILCs and induced by IL-33 and TSLP – in the regulation of ILC2-mediated inflammatory responses. RBM3 dampens both type 2 and 17 cytokine production by ILCs as well as lung inflammation in the setting of fungal allergen and IL-33 exposure. Transcriptomic analysis revealed RBM3 regulation of multiple cytokines, receptors, transcription factors, and survival genes critical to ILC function. These studies also highlight RNA-binding proteins as novel post-transcriptional regulatory mediators of ILC activation.

## Methods

### Mice

6-12-week-old female and male C57BL/6J mice were obtained from Jackson Laboratories (Bar Harbor, ME). Wild-type mice were age and gender matched to *rbm3*−/− mice acquired from Dr. Peter Vanderklish at Scripps Research and bred in house. *tslpr*−/− mice were acquired from Dr. Michael Croft at the La Jolla Institute for Immunology and originally from Dr. Steven Ziegler^67^. The *rbm3*−/−*rag2*−/− mice were created through multiple crosses of *rbm3*−/− and *rag2*−/− mice and bred in house. All studies were approved by the University of California, San Diego Institutional Animal Care and Use Committee.

### Lung inflammation models

Mice were intranasally challenged with *Alternaria alternata* extract (Greer, Lenoir, NC) or recombinant IL-33 (R&D Systems, Minneapolis, MN) diluted in PBS. For the isolation of the ILC subsets for RNA-seq analysis and the isolation of Lin-Thy1.2+ ILCs for *in vitro* assays, mice were challenged with 50μg *Alternaria* 4 times over 10 days. Mice were challenged with 25μg *Alternaria* 3 times over 7 days to expand the Lin-Thy1.2+ ILC population for RNA-seq analysis. Experiments with *tslpr*−/− mice or the anti-IL-33R blocking antibody involved 4 challenges with *Alternaria*. Various *Alternaria* challenged models were utilized with the *rbm3*−/− mice. Mice were intranasally challenged 3 times over 7 days with 10μg or 25μg *Alternaria* or were challenged once with 20μg *Alternaria* followed by 3 challenges of 10μg *Alternaria* over 10 days where indicated. WT and *rbm3*−/− mice were also intranasally challenged with 10ng recombinant IL-33 3 times over 7 days. *rag2*−/− and *rbm3*−/−*rag2*−/− mice were challenged with 20 μg *Alternaria* 4 times over 10 days. Wild-type mice were treated with anti-IL-33 antibody (DJ8)^28^ or control IgG intraperitoneally on Days −1, 0, 3, and 6.

### BAL and Lung Processing

Bronchoalveolar lavage (BAL) was collected in 2% BSA (Sigma, St. Louis, MO) and supernatant for the first flush of BAL was saved and stored in −20C for ELISA analysis^23, 28^. Lung was collected in RPMI and was dissociated into single-cell suspension using the Miltenyi Lung Digest Kit and Dissociator (Miltenyi Biotec, Bergisch Gladbach, Germany) as per the company’s protocol. Cells were counted using flow cytometry using the Novocyte cytometer (ACEA, San Diego, CA).

### Histology

In some experiments, the left half of the lung was used for histology. Lungs were perfused and fixed in 4% paraformaldehyde. Hematoxylin and Eosin (H&E) and Periodic acid–Schiff (PAS) staining was performed at the Histology Core in UCSD’s Moore’s Cancer Center and imaged with microscopy as previously reported^23^.

### ELISA

Samples stored at −20C were analyzed using IL-5 and IL-13 ELISA kits (R&D Systems, Minneapolis, MN) as per the company’s instructions. Plates were read using a microplate reader model 680 (Bio-Rad Laboratories, Hercules, CA). ELISA data was analyzed using Excel and Graphpad Prism (San Diego, CA).

### ILC purification and RNA Sequencing

FACS purification and cell preparation for RNA-seq analysis of the four ILC subsets was completed as previously described (GEO accession #136156)^23^. WT and *rbm3*−/− ILCs were sorted with the BD FACSAria II or the BD FACSAria Fusion at the UCSD Human Embryonic Stem Cell Core Facility. Lin-Thy1.2+ ILCs were sorted directly into TrizolLS. RNA-sequencing was performed at the La Jolla Institute.

Briefly, total RNA was purified using miRNAeasy kit (Qiagen), quantified and quality of RNA assessed by Fragment Analyzer (Advance Analytical)^68^. All samples had an RNA integrity number (RIN) > 9.0 and passed our quality and quantity control steps as described previously. Purified total RNA (≈5 ng) was amplified following the Smart-seq2 protocol^68, 69^. purified total RNA (≈5 ng) was amplified following the Smart-seq2 protocol^68, 69^. Briefly, mRNA was captured using poly-dT oligos and directly reverse-transcribed into full-length cDNA using the described template-switching oligo^68, 69^. cDNA was amplified by PCR for 15 cycles and purified using AMPure XP magnetic bead (0.9:1 (vol:vol) ratio, Beckman Coulter). From this step, for each sample, 1 ng of cDNA was used to prepare a standard NextEra XT sequencing library (NextEra XT DNA library prep kit and index kits; Illumina). Barcoded Illumina sequencing libraries (NextEra; Illumina) were generated utilizing an automated platform (Biomek FXP, Beckman Coulter). Both whole-transcriptome amplification and sequencing library preparations were performed in a 96-well format to reduce assay-to-assay variability. Quality control steps were included to determine total RNA quality and quantity, the optimal number of PCR preamplification cycles, and fragment library size. The reference genome was mm10 (mouse genome). None of the samples failed quality controls and were pooled at equimolar concentration, loaded and sequenced on the Illumina Sequencing platform, Novaseq 6000 (Illumina). Libraries were sequenced to obtain more than 20 million 100 × 100 bp paired-end reads (S4 P200 flow cell and sequencing Kit; Illumina) mapping uniquely to mouse mm10 mRNA reference.

Sequencing data for this study has been deposited into the Gene Expression Omnibus under accession #136156. The paired-end reads that passed Illumina filters were filtered for reads aligning to tRNA, rRNA, adapter sequences, and spike-in controls. The reads were then aligned to mm10 reference genome using STAR(v 2.6.1c)^70^. DUST scores were calculated with PRINSEQ Lite (v 0.20.3)^71^ and low-complexity reads (DUST > 4) were removed from the BAM files. The alignment results were parsed via the SAMtools^72^ to generate SAM files. Read counts to each genomic feature were obtained with the featureCounts program (v1.6.5)^73^. After removing absent features (zero counts in all samples), the raw counts were then imported to R/Bioconductor package DESeq2^74^ to identify differentially expressed genes among samples. P-values for differential expression are calculated using the Wald test for differences between the base means of two conditions. These p-values are then adjusted for multiple test correction using Benjamini Hochberg algorithm^75^ to control the false discovery rate.

### qPCR

RNA was reverse transcribed to cDNA using a Transcriptor First Strand cDNA Synthesis Kit (Roche) according to the manufacturer’s instructions. RT-PCR was performed using SYBR Green I Master (Roche) and RT-PCR–specific primers. The primer sequences (5′–3′) were as follows: mIL-5 forward AAGAGAAGTGTGGCGAGGAGA; mIL-5 reverse CACCAAGGAACTCTTGCAGGTAA; mIL-13 forward GAGCAACATCACACAAGACCAGA; mIL-13 reverse GCCAGGTCCACACTCCATA; CysLT1R forward AACGAACTATCCACCTTCACC; CysLT1R reverse AGCCTTCTCCTAAAGTTTCCAC; L32 forward GAAACTGGCGGAAACCCA; and L32 reverse GGATCTGGCCCTTGAACCTT. qPCR was completed using the Rbm3 transcript variant 2 (NM_001166409). Transcripts were measured relative to L32 using Roche LightCycler 480 (Roche Diagnostics)^64^.

### Flow Cytometry

For surface stains, one million lung and BAL cells were stained. For intracellular stains, two million lung cells were stained. Fc receptors were first blocked for 5 minutes using CD16/CD32 (Biolegend, San Diego, CA). Eosinophils were identified as CD11c-Siglec-F+ and neutrophils were identified as SiglecF-GR-1+; they were stained using CD45.2 (PerCP), Siglec-F (PE), GR-1(APC), and CD11c (FITC). ILCs were identified as Lineage-Thy1.2+ lymphocytes or Lineage-T1ST2+ lymphocytes and were stained using CD45.2 (PerCP), Thy1.2 (APC), T1ST2 (PE), and a lineage cocktail. The lineage cocktail (FITC) consisted of the BioLegend Lineage cocktail (consists of CD3e, Ly-6G/Ly-6C, CD11b, CD45R/B220, and TER-119), CD11c, NK1.1, CD5, FcεR1, TCR*β*, and TCRγδ. The ILC subsets were stained using ST2 (APC) and CD127 (PE-Cy7). For nuclear intracellular staining, cells were permeabilized using the FoxP3 kit (ThermoFisher, Waltham, MA) after surface staining. Cells were stained using Ki-67 (PE or APC), RBM3, GATA3 (PE), and ID2 (PE).

For cytokine intracellular staining with the 10-day challenge model, cells were cultured overnight with Golgi Plug (Fisher Scientific, Hampton, NH) at 500,000 cells per well. After surface staining for ILCs, cells were fixed and permeabilized using the BD kit (BD Biosciences, La Jolla, CA) and stained for IL-5 (PE) or IL-13 (PE). For cytokine intracellular staining following the 7-day IL-33 and *Alternaria* challenge model, lung cells were cultured for 3 hours with cell stimulation cocktail (ThermoFisher, Waltham, MA) at 1 million cells per well. After surface staining for ILCs, cells were fixed and permeabilized using the BD kit and stained for IL-5 (PE), IL-13 (PE), or IL17A (eFlour506). Lung cells stained for Bcl-2 expression were surfaced stained for ILCs, fixed and permeabilized with the BD kit, and stained with Bcl-2 (PE-Cy7).

For human PBMC and nasal polyp staining, ILC2s were identified as Lineage-, CRTH2+ lymphocytes. The lineage cocktail (FITC) consisted of CD3, CD14, CD16, CD19, CD20, CD56, TCRγδ, CD4, CD11b, CD235a, and FcεRI. Cells were permeabilized using the FoxP3 kit and stained for RBM3. The polyclonal RBM3 antibody used in this study was raised in rabbits to the 14 c-terminal amino acids of RBM3, and affinity purified to the immunizing peptide. As described in prior work^76, 77^, the affinity purified anti-RBM3 antibody recognizes an ~17 kDa band corresponding to RBM3 on Western blots and selectively labels RBM3 in situ under a variety of fixation conditions. Flow Cytometry was performed using the Novocyte. Data was analyzed using FlowJo software (Tree Star, Ashland, OR). All antibodies were obtained from Biolegend, ThermoFisher, or BD Biosciences.

### Immunofluorescence

For mouse airways, immunofluorescence for RBM3 was performed on naïve and *Alternaria* challenged airways as previously reported^78, 79^. Cell nuclei were stained with DAPI. Images were taken from at least 5 airways of at least 3 mice per group.

### ILC and PBMC cultures

WT and *rbm3*−/− Lin-Thy1.2+ ILCs were sorted using the BD FACSAria Fusion and BD FACSAria II sorters from UCSD’s Human Embryonic Stem Cell Core Facility. Collected ILCs were rested in 10ng/mL IL-2 and IL-7 (R&D Systems, Minneapolis, MN) in T-cell media (TCM) for 48 hours. ILCs were cultured in a 96-well plate at 40,000 cells per well. Pre-stimulation media was collected and stored in −80C for ELISA. ILCs were stimulated for 24 hours with 15ng/mL or 30ng/mL IL-33 (R&D Systems, Minneapolis, MN) in TCM. Post-stimulation supernatant was collected and stored at −80C for ELISA analysis.

Human peripheral blood ILC2s were sorted as CD45+lin-CRTH2+ lymphocytes and cultured and treated with TSLP and IL-33 before being fixed with 4% PFA and processed for immunocytochemistry using a RBM3 antibody (at 1:2000) and a Cy3 secondary. Cells were also stained for DAPI. Images were taken at 20X. Immuno-positive elements were captured and analyzed by thresholding the intensity histogram in the Cy3 channel at 100. Data for objects of 50-175 pixels were included. In some studies, human PBMCs were collected and cultured overnight with 10ng/mL IL-2 and stimulated with 10ng/mL IL-33, 50ng/mL TSLP, or a combination of both and RBM3 analyzed by intracellular flow cytometry.

### Human samples

Human Subjects Committee approval was granted at UC San Diego for all studies involving human nasal and blood samples.

### Statistical Analysis

Statistical analysis was performed with GraphPad Prism software (GraphPad Software, La Jolla, CA). P-values were obtained using the Mann-Whitney test or the unpaired t-test and a P value of less than 0.05 was considered statistically significant such that * p < 0.05, ** p < 0.01, *** p < 0.001.

---

**Supplementary Fig. 1:**
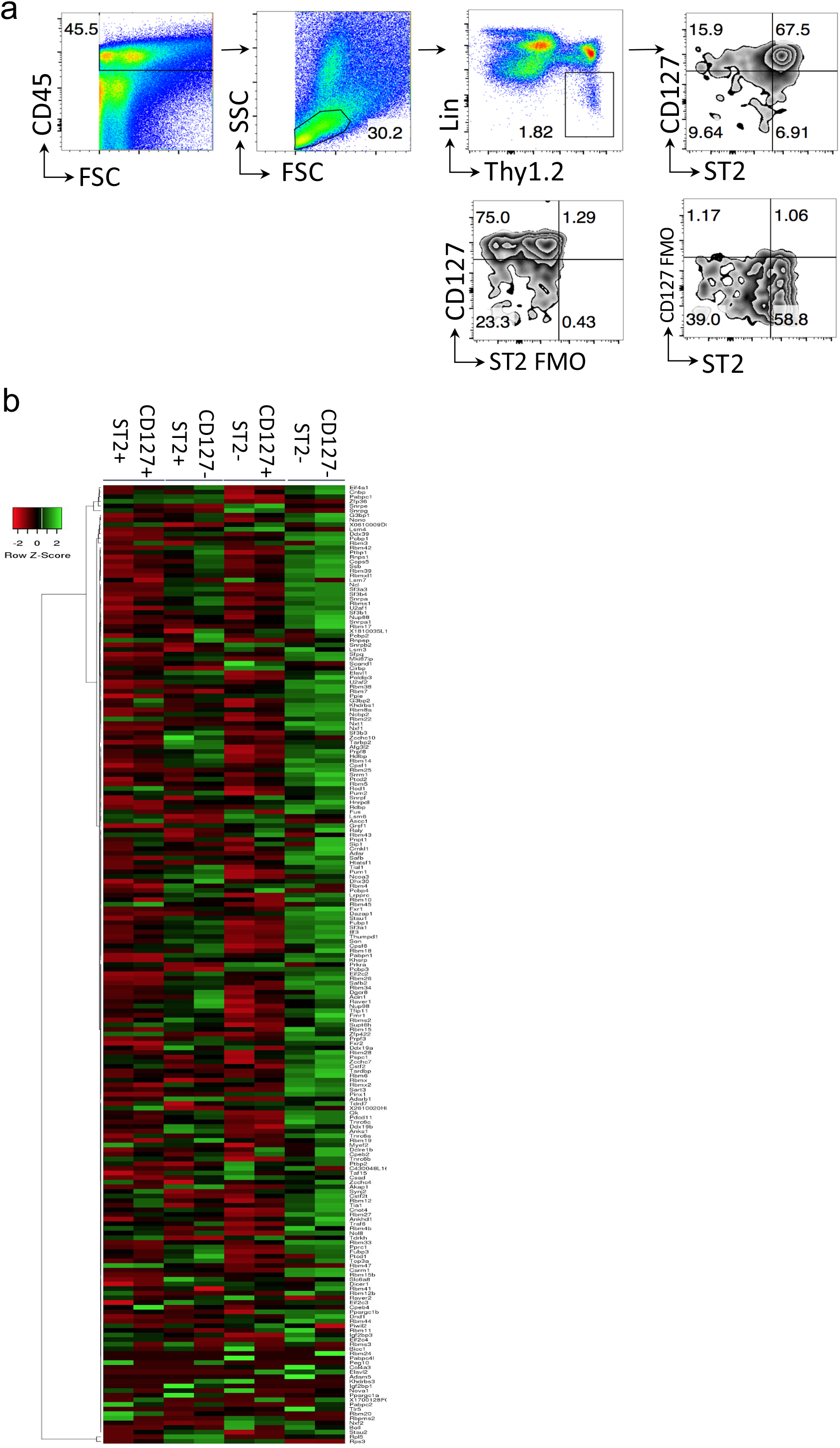
Gating strategy for ILC subsets collected for bulk RNA-Seq analysis and RBM transcriptome. (a) ILCs were gated as CD45+ Lineage-Thy1.2+ lymphocytes. Subsets were identified by their expression of ST2 and CD127. Gating was based on FMOs. (b) Wild-type mice were challenged 4 times with 50μg *Alternaria* over the span of 10 days and four ILC subsets (CD127+ST2, CD127-ST2+, CD127+ST2-, CD127-ST2-) were sorted and collected for RNA-seq. Heatmap showing relative expression of 207 RBPs in the four sorted ILC subtypes using Centroid linkage and Manhattan clustering

**Supplementary Fig. 2:**
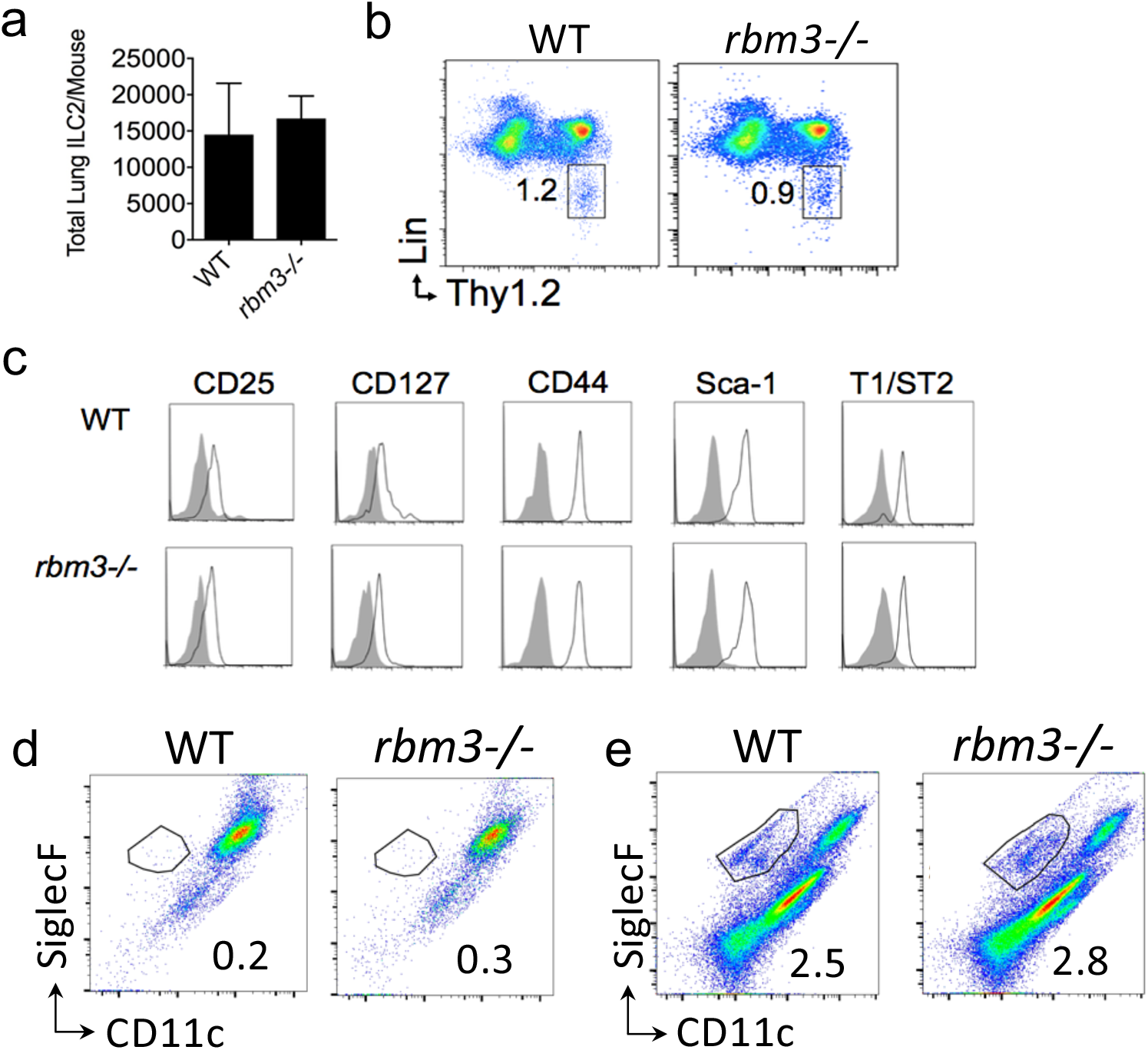
Naive rbm3−/− mice have similar lung ILC numbers, surface marker expression, and eosinophilia compared with WT mice. Naive wild-type and *rbm3*−/− lungs were stained for ILCs and eosinophils. (a) Total lung ILCs. (b) Representative FACS plots of Lin-Thy1.2+ ILCs. (c) Histograms of surface marker expression on naive lung ILCs. Grey = Isotype control. Representative FACS plots of naïve (d) BAL eosinophils and (e) lung eosinophils.

**Supplementary Fig. 3:**
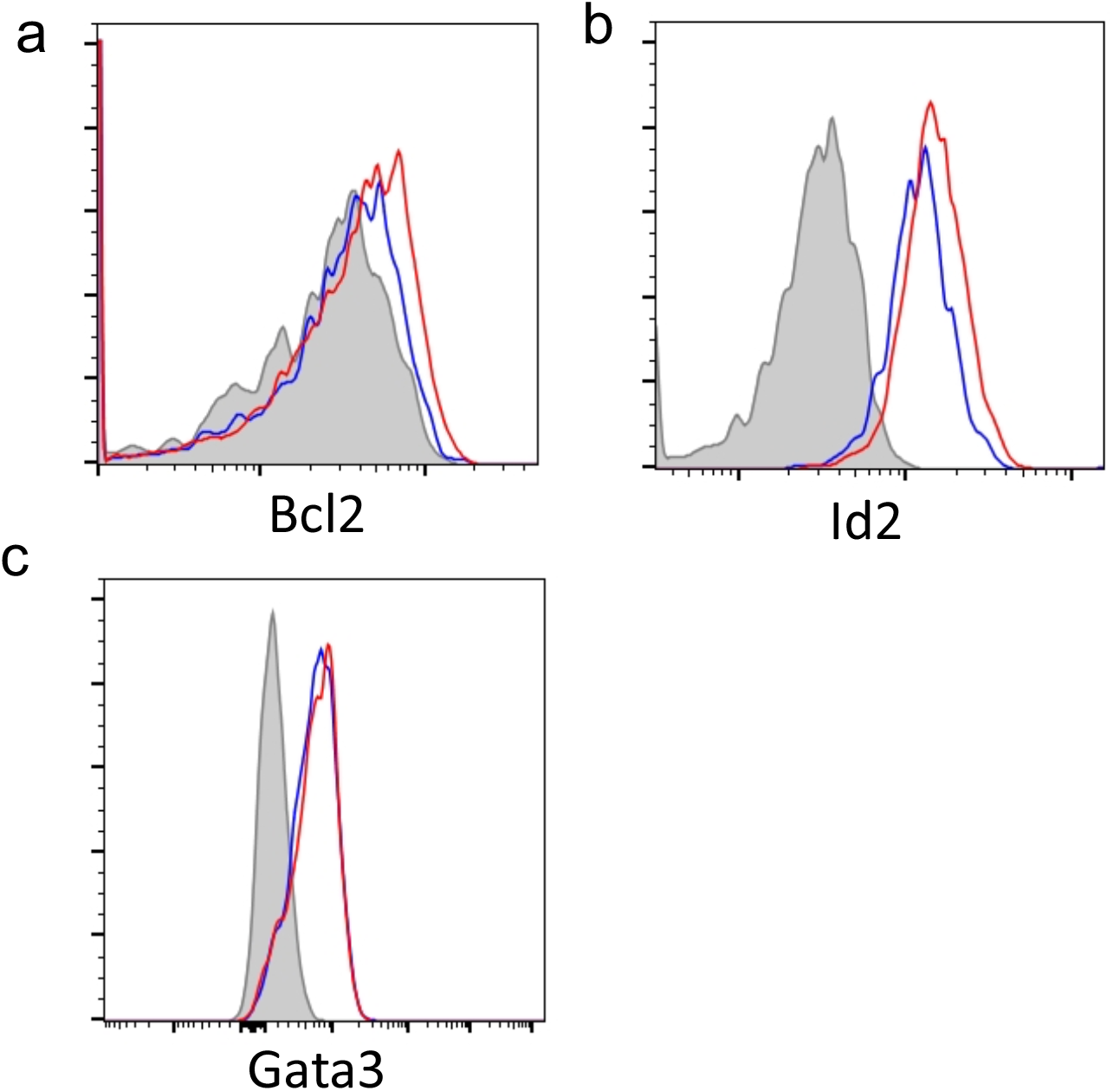
Expression of Bcl2, Id2, and GATA3 by flow cytometry in WT and rbm3−/− mice. WT and *rbm3*−/− mice were challenged with 25 μg *Alternaria* three times over 7 days. Data representative of 4 mice per group. (a) Histogram plot of ILC Bcl2 expression. (b) Histogram plot of ILC Id2 expression. (c) Histogram plot of ILC GATA3 expression. Blue = WT. Red = *rbm3*−/−. Grey = isotype control.

**Supplementary Fig. 4:**
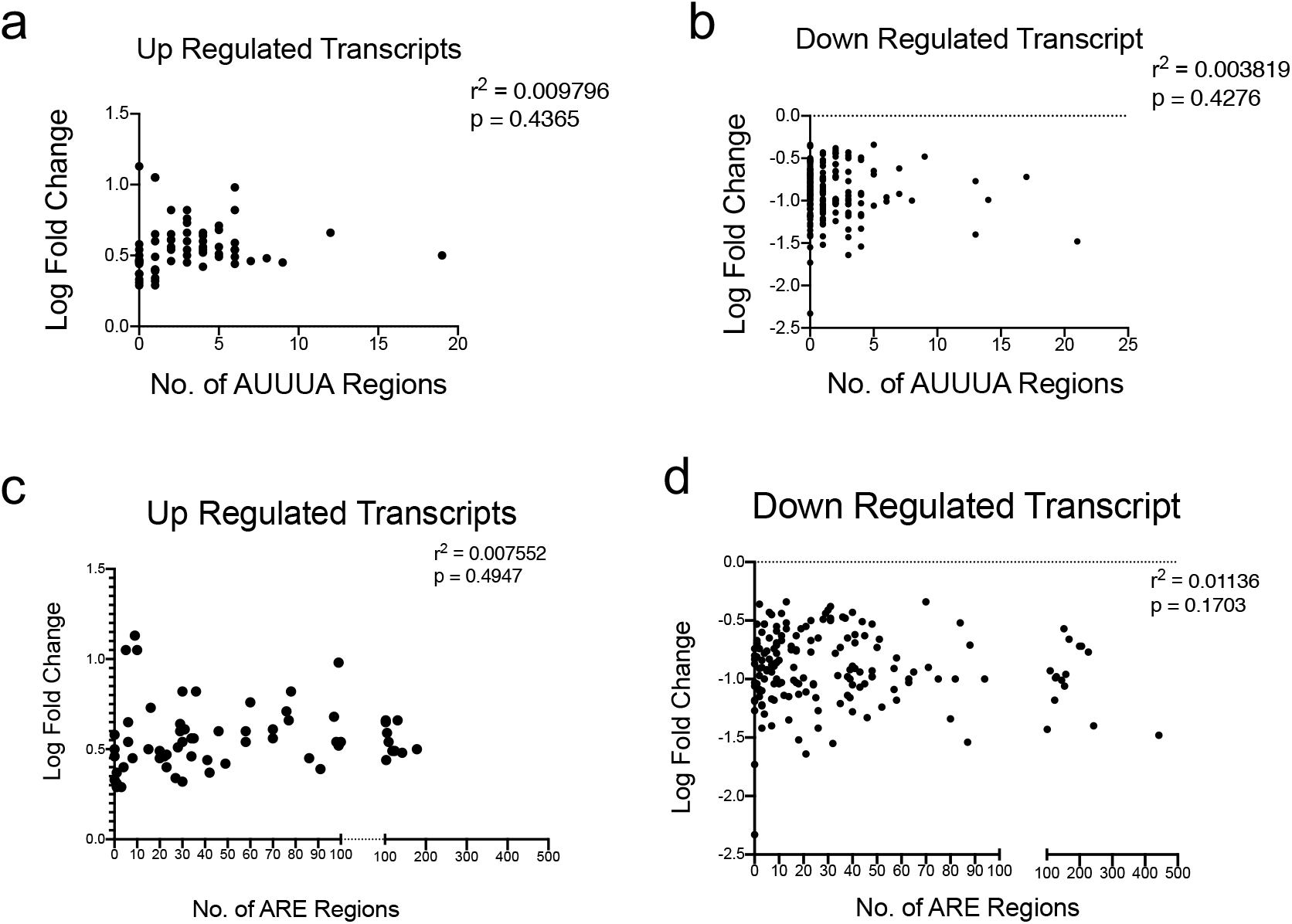
Lack of correlation between transcript AU-rich element (ARE) numbers with rbm3−/− ILC transcriptome. WT and *rbm3*−/− mice were challenged with 25 μg *Alternaria* three times over 7 days. Lin-Thy1.2+ ILCs were FACS-sorted and bulk RNA-sequenced. Comparison of (a) up-regulated and (b) down-regulated transcripts to their number of AUUUA regions using AREsite http://rna.tbi.univie.ac.at/AREsite. Comparison of (c) up-regulated and (d) down-regulated transcripts to their total number of ARE regions using AREsite http://rna.tbi.univie.ac.at/AREsite.

